# When does additional information improve accuracy of RNA secondary structure prediction?

**DOI:** 10.1101/2025.03.18.643972

**Authors:** Logan Rose, Luis Sanchez Giraldo, Duc Nguyen, Matthew Wheeler, David Murrugarra

**Affiliations:** Department of Mathematics, University of Kentucky, Lexington, KY 40506, USA; Department of Electrical and Computer Engineering, University of Kentucky, Lexington, KY 40506, USA; Department of Mathematics, University of Tennessee, Knoxville, TN 37996, USA; Department of Medicine, University of Florida, Gainesville, FL 32610, USA

## Abstract

The secondary structure of an RNA sequence plays an important role in determining its function, and accurate prediction of the structure is still a major goal in computational biology. Improvements in the prediction accuracy of the secondary structure can be achieved via auxiliary information. In this paper, we study features based on suboptimal formations competing with the minimum-free energy formation and investigate their role in determining the improvement of accuracy via auxiliary information, which we call directability. Here, we introduce a similarity measure among competing substructures called profiles. Then, we present an *n*-dimensional representation of the profiles which allows the use of topological data analysis (i.e., persistence landscapes) to obtain different metrics that represent topological features. Then, we built random forest classifiers using these novel features. We show how the similarity feature is more important for classifiers trained on sequences with similar structures while the topological features are more important for classifiers trained on sequences with dissimilar structures. We perform extensive testing on two sets of RNA sequences where we studied the sensitivity of the classification accuracy and their feature importance.

## 1 Introduction

An RNA chain folds back to form higher dimensional structures such as patterns of base pairings forming helices and single stranded loops. The two-dimensional formation is called the *secondary structure of an RNA sequence*. Figure 1 shows the secondary structure of *tRNA* which was generated using the software VARNA [1]. The secondary structure plays an important role in determining the function of an RNA sequence, where a mutated structure might be involved in several diseases [2, 3].

**Figure 1:**
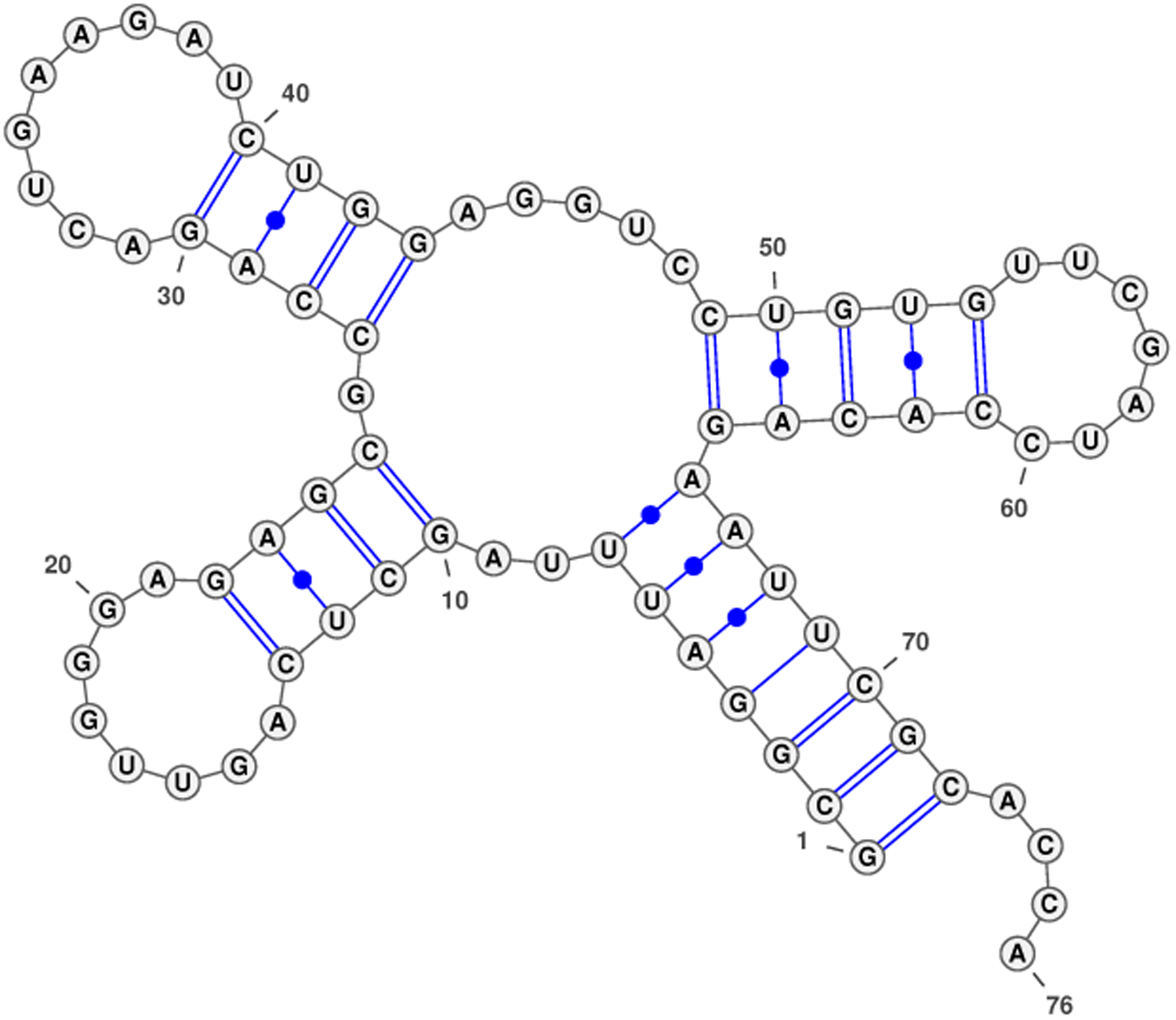
*tRNA* secondary structure which was generated using VARNA [1].

One of the most common tools for computational prediction relies on the nearest neighbor thermodynamic model (NNTM) [4], which is based on energy minimization using dynamic programming. There are several implementations of NNTM including *RNAstructure* [5], the *ViennaRNA* Package [6], *UNAFold* [7], and *GTfold* [8]. Although very useful, the physics-rooted NNTM approach is prone to accuracy errors in its predictions [9, 10]. Accuracy can be improved by incorporating additional information, such as high-throughput chemical probing data that further constrain the thermodynamic model [11, 12, 13]. However, providing such additional information directly requires costly procedures [11]. Alternative methods that predict such auxiliary information using machine learning (ML) and deep learning (DL) have been proposed [14, 15, 16]. In recent years, data-driven methods for RNA structure prediction based on machine learning (particularly deep learning) have also been proposed [3, 17, 16, 18]. These methods are based on known sequence to secondary structure maps to learn a model to predict the secondary structure of unseen sequences.

The newer methods based on ML, especially deep learning, for RNA structure prediction seem to provide higher accuracies but give little insight about the mechanisms for folding. These methods may be able to learn nuances missed by the NNTM, but rely heavily on training data. Therefore, the imbalance of RNA families in training data sets and the fact that different RNA families have different patterns can create prediction issues, such as overfitting from deep learning methods, where a method performs well in one RNA family and underperforms in others [15, 16]. Addressing these challenges will require going beyond the single minimal free energy prediction, such as studying the suboptimal structure formations to understand the competing alternatives. Boltzmann sampling of secondary structures [19] allows the study of the competing substructural alternatives. RNA structure profiling [2] is a tool based on graph theory techniques that allows the representation of the competing substructures as a partially ordered set that highlights the competing substructures in a Boltzmann sample. Many of these tools have not been fully exploited yet for predicting the secondary structure of RNA.

In this paper, we investigate topological features from the graph of competing substructures that are relevant for determining whether a sequence can significantly improve its predicted accuracy using synthetic auxiliary data. To this end, we construct classifiers that take these features as input to predict 1) directability (i.e., improvements of accuracy via auxiliary information) and 2) well-prediction (that is, if the accuracy of the structure prediction will be above certain threshold). The rest of the paper is structured as follows. In the Background section, we describe the RNA structure profiling approach. In the Methods section, we introduce our similarity measure among profiles and describe an *n*-dimensional embedding for these profiles to perform topological data analysis. In the Results section, we report the accuracy and feature importance of different classifiers for directability and well-prediction. Finally, we describe the main takeaways of the paper in the Conclusions section.

## 2 Background

### 2.1 RNA Structure Profile graphs

For an RNA sequence, we obtain features based on suboptimal formations competing with the minimum-free energy (MFE) formation using RNA structure profiling. RNA structure profiling [2] is a tool for understanding competing structural alternatives based on Boltzmann samplings [19]. RNA profiling provides a summary graph as a poset (i.e., a partially ordered set) that highlights the competing substructures from a Boltzmann sample. Figure 2 shows a profile graph for *tRNA*. The table in the top left provides the information of the selected profiles (those with frequency above a certain threshold). Boxed vertices indicate selected profiles, while dashed ovals indicate the intersection of two profiles. Each node is labeled with the profile in parenthetic notation along with its specific and general frequencies written as a ratio.

**Figure 2:**
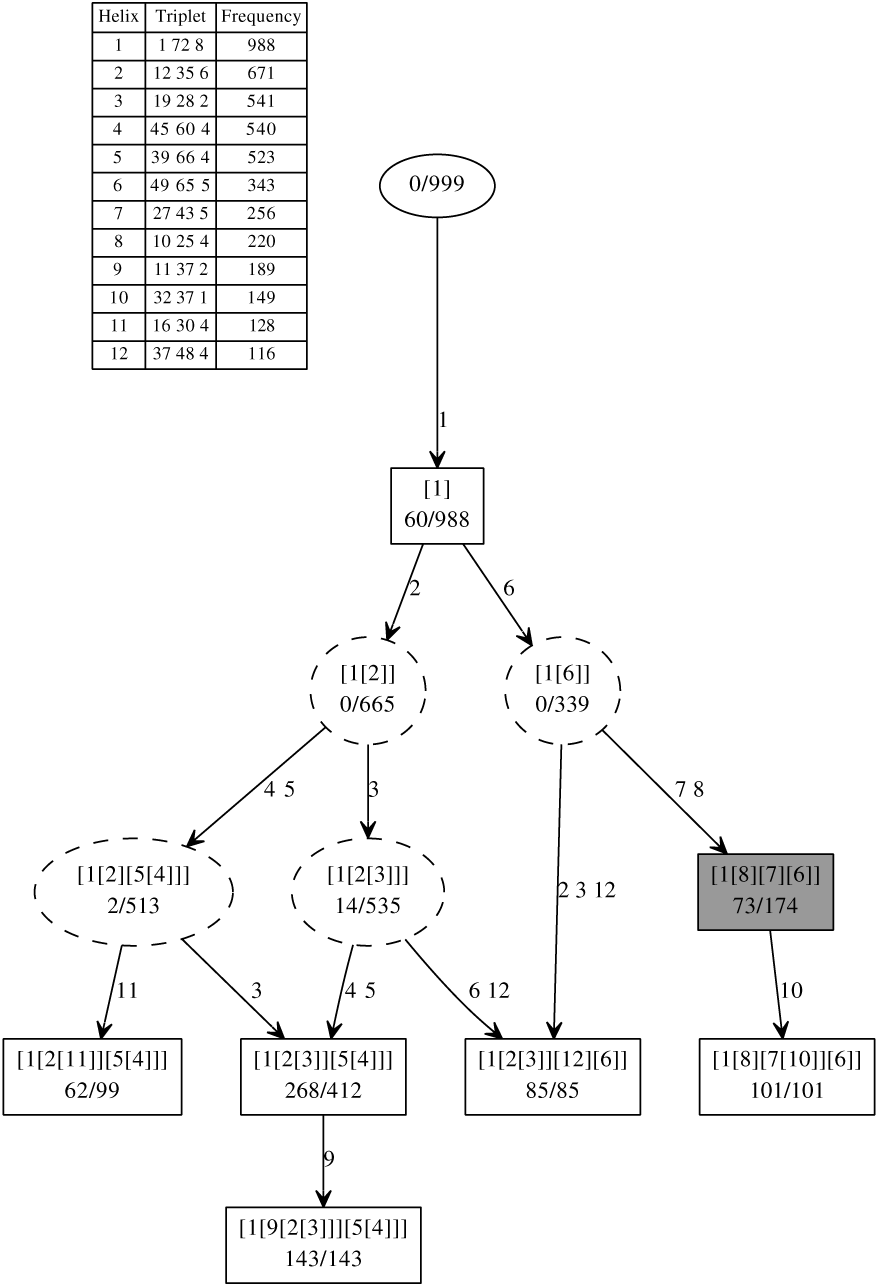
Profile graph for *tRNA*. The table in the top left provides the information of the selected profiles. The shaded profile is the native profile corresponding to the correct structure. Figure generated using RNA Structure Profiling [2].

The profiling method uses the triplet notation of a helix. For instance, the first helix *h* = (1, 72, 8) in the table of Figure 2 has 8 consecutive base pairs {(1, 72), (2, 71), (3, 70), (4, 69), (5, 68), (6, 67), (7, 66), (8, 65)}. In general, consecutive base pairs {(*i, j*), *…,* (*i* + *k* − 1, *j* − *k* + 1)} form a helix *h* = (*i, j, k*). A helix (*i, j, k*) is maximal if (*i* − 1, *j* + 1) and (*i* + *k, j* − *k*) would be non-canonical base pairs or violate *j* − *i* − 2*k* ≥ 2. Two helices *x* and *y* are related if their base pairs are a subset of the same maximal helix *h*. The equivalence class under this relation is called a helix class, denoted by [*h*]. Given a Boltzmann sample S, the frequency of [*h*] is freq([*h*]) = {*s* ∈ S : [*h*] ∩ *s* ≠ ∅}*/*|S|. Profiling groups together similar substructures using helix classes.

The profile *p_s_* of a structure *s* is the set of its corresponding helix classes that can be presented in parenthetic notation. For instance, the shaded profile in Figure 2 is composed of four helix classes and is represented by [1[8][7][6]]. The set of base pairings (in triplet notation) of the composing helix classes are provided in the table at the top left of Fig. 2.

### 2.2 Topological Data Analysis

With the continual increase in size and complexity of data sets, building analytical tools for understanding this complexity is important. Many of these complex data sets admit interesting topological and geometric features. For low-dimensional data, this can many times be inferred by inspection, but for high-dimensional data, more advanced topological tools are required. Topological data analysis focuses on the development and implementation of computational tools built on ideas from algebraic topology to identify and quantify these features. One particularly versatile tool is that of Persistent homology which ascribes a filtration of simplicial complexes *K*_•_ to a given data set,

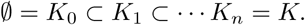

There are a number of ways to construct these filtrations from point cloud data, and in what follows we will use the Vietoris-Rips filtration [20]. Given a filtration, one then computes the homology at each step of the filtration which induces a morphism of groups for each inclusion map of the filtration. The resulting persistence module is a sequence of group morphisms

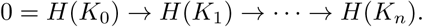

Any topological feature in this persistence module is born at some filtration level *b* and then dies at a subsequent filtration level *d*. Usually this topological information can be encoded as a persistence diagram, a collection of points *D* = ⊕*I_i_* ⊂ {(*x, y*) ∈ ℝ^2^| *x < y*}, representing these topological features where each tuple *I_i_* = (*b_i_, d_i_*) corresponds to the birth and death of a feature [21]. However in order to perform statistical analyses, a more appropriate topological summary for our purposes is the persistence landscape [22, 23]. Given a persistence diagram *D*, the persistence landscape can be defined as follows. Consider the functions

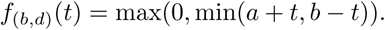

The persistence landscape is then given by the sequence of functions (*λ*_1_, *λ*_2_, *…*) where

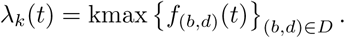

The space of persistence landscapes forms a Hilbert space with norm induced by the inner product below

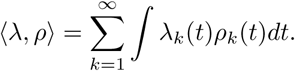

Persistence landscapes have a number of nice properties that make them ideal for statistical analysis. First, they are stable with respect to the original point cloud, that is, small perturbations of the data result in only small perturbations of the corresponding landscapes. Being a Hilbert space, persistence landscapes also come with operations of addition and scalar multiplication allowing one to calculate averages, standard deviations, and variances of persistence landscapes as opposed to persistence diagrams.

### 2.3 Embedding profile graphs into an *n*-dimensional hypercube

In order to apply topological data analysis tools to extract features based on the the properties of the profile graph, we map the vertices of the profile graph *G* to points of an *n*-dimensional hypercube *D*. That is, we map each vertex of *G* to a point in *D* as follows. First, we list all helix classes *h*_1_, *…, h_n_*in the Boltzmann sample. Then, we use the ratio of the specific frequency of the profiles to the frequency of the composing helix classes to represent the profiles as points in ℝ*^n^*. More specifically, we use the binary random variable 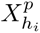 that indicates the presence of the helix class *h_i_* in profile *p*. Then, we define the coordinates of the corresponding point as follows. Let

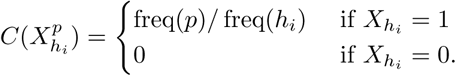

Finally, each profile with *p* is mapped to the point 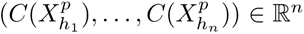. Figure 3 shows the profile graph for *Adenine* and its hypercube representation. This profile graph has only 1 node and the corresponding point is shown in red in Figure 3. We use this hypercube representation to get novel topological features for the classifier of RNA sequence directability.

**Figure 3:**
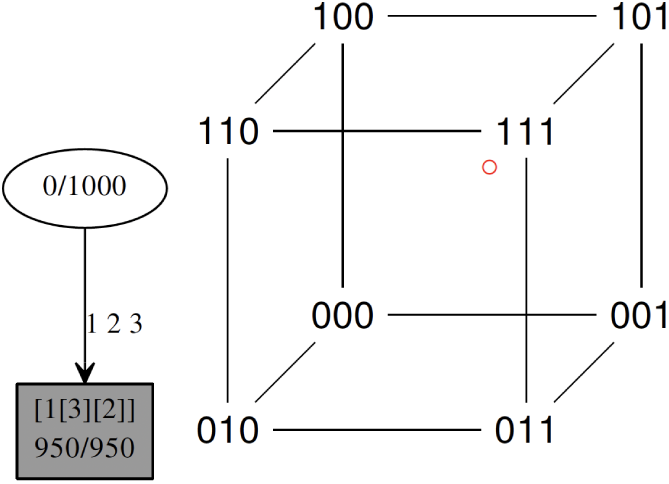
Profile graph of a boltzmann sample for *Adenine* and its hypercube representation. The shaded profile is the native profile. The red dot represents the point: 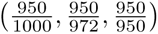.

## 3 Methods

Here we define the similarity measure among profiles and the norms of persistence landscapes that will represent topological features of the profile graph. Then, we will use these metrics to extract relevant features for our directability and well-prediction classifiers.

### 3.1 Similarity and variance measures for profiles

For RNA structure prediction, the standard measures for assessing the accuracy of the predictions are the positive predictive value (PPV), 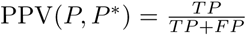, and the sensitivity (SEN), 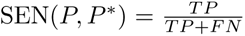, where

- *TP*: true positives. Base pairings that occur in both the native and the predicted structure.
- *FP*: false positives. Base pairings that occur in the predicted structure but not the native.
- *FN*: false negatives. Base pairings that occur in the native structure but not the predicted.
- *PPV*: is the fraction of true positives in the predicted structure.
- *SEN*: is the fraction of true positives in the native structure.

While SEN and PPV have been primarily used as standard measures for structures, they can also be applied for RNA profiles in the following way. Given a profile *P* in a Boltzmann sample, we take the MFE profile *P* ^∗^ as a reference and compare *P*to *P* ^∗^ using,

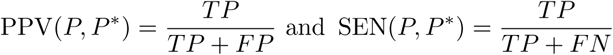

where

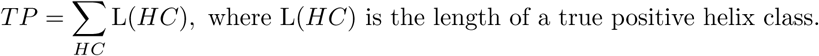

The summation is over all true positive helix classes. *FP* and *FN* are defined in a similar way. L(*HC*) could also be defined as the average length of helices in the helix class. We take the MFE profile as a reference because we want to measure how centered is the sample in the MFE structure.

For each profile *P_i_*, we compute PPV(*P_i_, P* ^∗^) and SEN(*P_i_, P* ^∗^). Then, we define the similarity of *P_i_* and *P* ^∗^ as the arithmetic mean of PPV(*P_i_, P* ^∗^) and SEN(*P_i_, P* ^∗^):

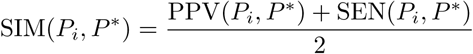

Since each profile comes with its frequency, we can compute the weighted average of the SIM(*P_i_, P* ^∗^)’s,

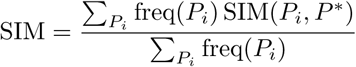

and the weighted sample SIM variance

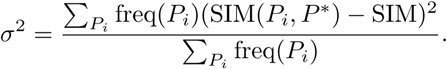

### 3.2 Persistence Landscapes

To detect topological features of the structure profile graph, we used persistence landscapes as a topological summary. For each profile graph, the corresponding embedding into the *n*-dimensional hypercube maps each profile *p* as

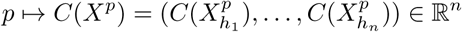

and thus the profile graph can be viewed as a point cloud in ℝ*^n^*.

To compute persistent homology, we constructed the Vietoris-Rips complex on this point cloud. For each *r >* 0, one associates the collection of simplices [*p*_1_, *…, p_n_*] satisfying *d*(*C*(*X^pi^*), *C*(*X^pj^*)) *< r* for all *p_i_, p_j_* in the simplex, where *d* here is the usual Euclidean metric. From there one computes homology of varying degree and subsequently the persistence landscapes.

All computations for calculating persistent homology and persistence landscapes were done using the tda-tools package in R. The landscapes were discretized by using an interval of 0.01 transforming each landscape into a high-dimensional vector. From there we computed the Euclidean norms of these landscapes. By taking homology of each degree *k*, we have a corresponding norm which we denote as the *k*-norm. We used the 0-norm and the 1-norm as our final summaries. The 0-norm describes the persistence of the connected components of the profile graph where the connected components correspond to each profile of the profile graph. The 1-norm then corresponds to the persistence of all 1-dimensional holes in the profile graph.

### 3.3 Synthetic auxiliary data

Experimentally derived auxiliary data such as Selective 2’-hydroxyl acylation analyzed by primer extension (SHAPE) has been used to improve accuracy. In this work, we use synthetically generated auxiliary data using the approach described in [15]. For an RNA sequence, the method in [15] produces a sequence *p* of the same length as the original RNA sequence, where *p_i_* is the predicted probability that the nucleotide in position *i* is paired. Using these predictions, one can convert each predicted probability *p_i_*to a SHAPE value to be associated with nucleotide *i*. Then, the following piecewise linear function is used to generate new synthetic auxiliary data:

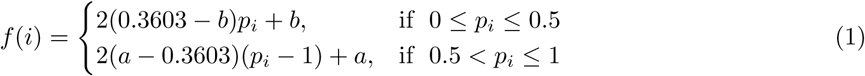

We list the main properties of this function below:

- This function has range [*a, b*],
- *f* (*i*) = *a* if *p_i_* = 1,
- *f* (*i*) = *b* if *p_i_* = 0, and
- *f* (*i*) = 0.3603 if *p_i_* = 0.5.
- To determine the values of *a* and *b*, we considered experimentally collected SHAPE data from two E. coli sequences: one 16S sequence and one 23S sequence from [12].
- We took the mean SHAPE value among both paired nucleotides and unpaired nucleotides; these values are 0.214 and 0.6624, respectively. Our experimental results used *a* = 0.214 and *b* = 0.6624.

For this paper, since our objective is to evaluate the role of the features, we assume to know the state of the nucleotide (as being paired or unpaired), and we generate our synthetic data with this information instead of using a predictor of the pairings.

### 3.4 Features

We considered the following features for the classifiers. Here, we provide the abbreviations that we will use in the figures in the Results Section:

1. Sim: the similarity before SHAPE.
2. Var: the variance before SHAPE.
3. SHS: the similarity after SHAPE.
4. VSH: the variance after SHAPE.
5. N0: the 0-norm before SHAPE.
6. N0SH: the 0-norm after SHAPE.
7. N1: the 1-norm before SHAPE.
8. N1SH: the 1-norm after SHAPE.

The first four features (i.e., Sim, Var, SHS, VSH) represent similarity features while the last four (i.e., N0, N0SH, N1, N1SH) represents topological features. The similarity features measure how similar the profiles are among themselves with respect to the MFE profile. The topological features measure the shape of the profile graphs. Intuitively, the 0-norm measures the number of nodes in the profile graph which correspond to the number of profiles available in the sample while the 1-norm describes the number of holes in the graph which correspond to the number of number of conflicts in suboptimal formations.

### 3.5 Test sequences

We have trained and tested several classifiers using known data sets of RNA sequences that have been used for other benchmarking purposes [24, 3]. We have used TrainSetA, TestSetA, TrainSetB, and TestSetB, which are available in the supplementary material of [3]. These data sets were originally established by Rivas et al. in [24]. As noted in [3], TrainSetB and TestSetB contain dissimilar structures, while TrainSetA and TestSetA have similar structures. The MFE accuracies before and after synthetic SHAPE for sets TrainSetA and TrainSetB are shown in Figures 4 and 5. Similarly, the MFE accuracies before and after synthetic SHAPE for TestSetA and TestSetB are shown in Figures 6 and 7. We obtained these accuracies using RNAStructure [5].

**Figure 4:**
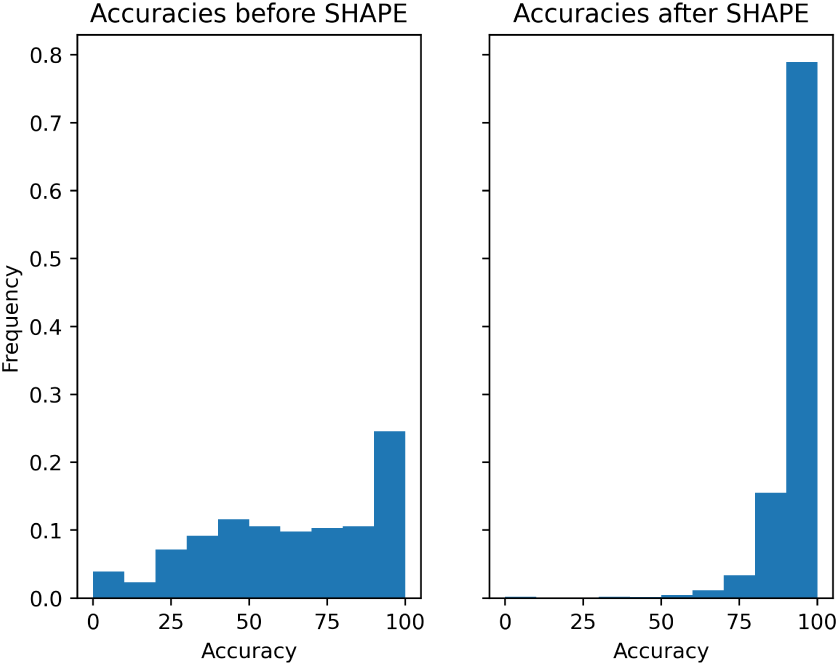
Accuracies before and after synthetic SHAPE for TrainSetA.

**Figure 5:**
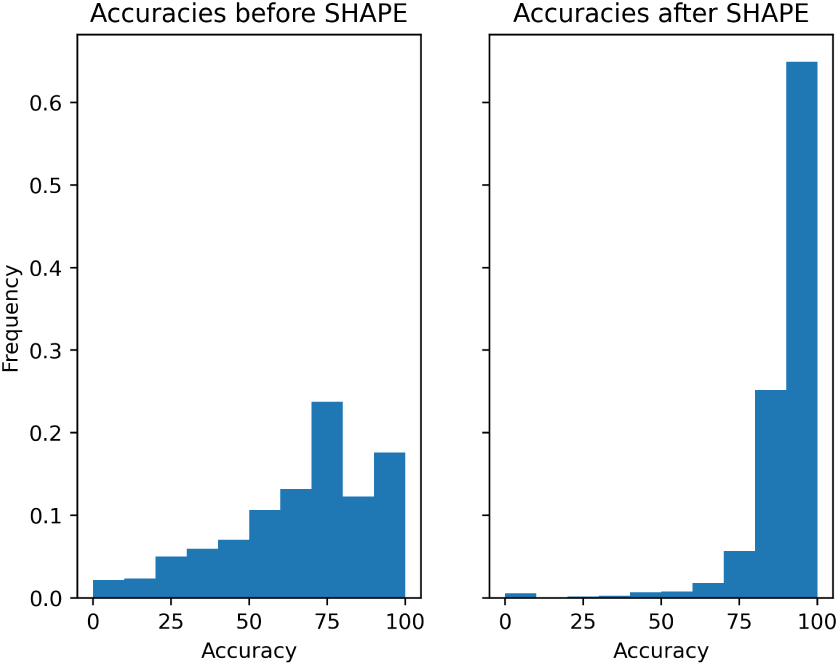
Accuracies before and after synthetic SHAPE for TrainSetB.

**Figure 6:**
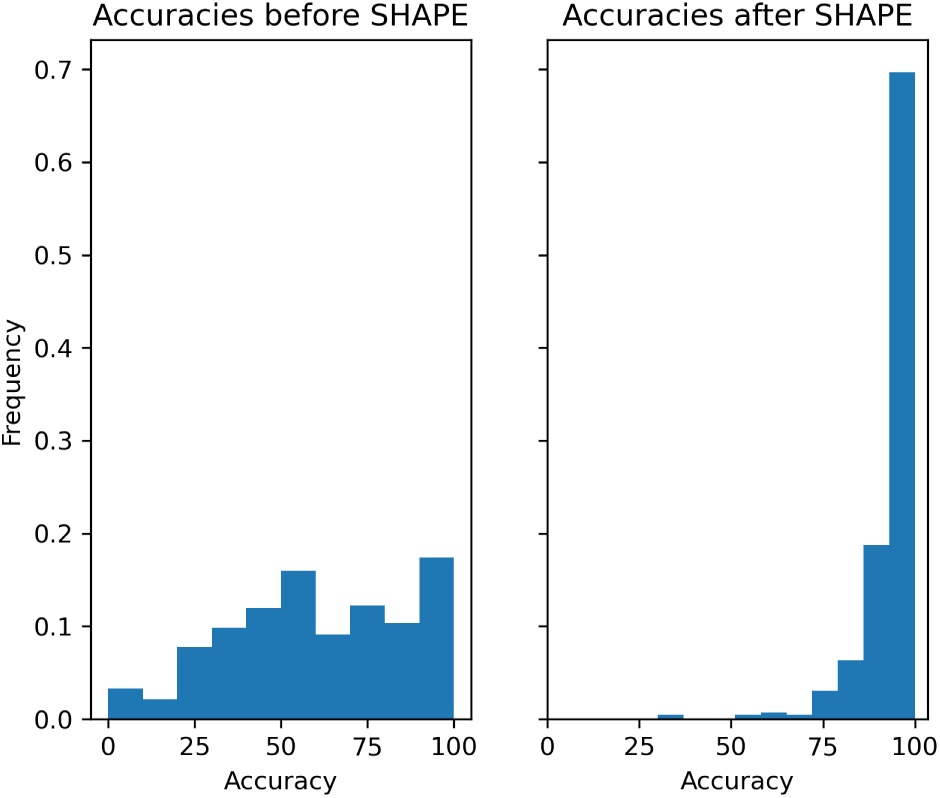
Accuracies before and after synthetic SHAPE for TestSetA.

**Figure 7:**
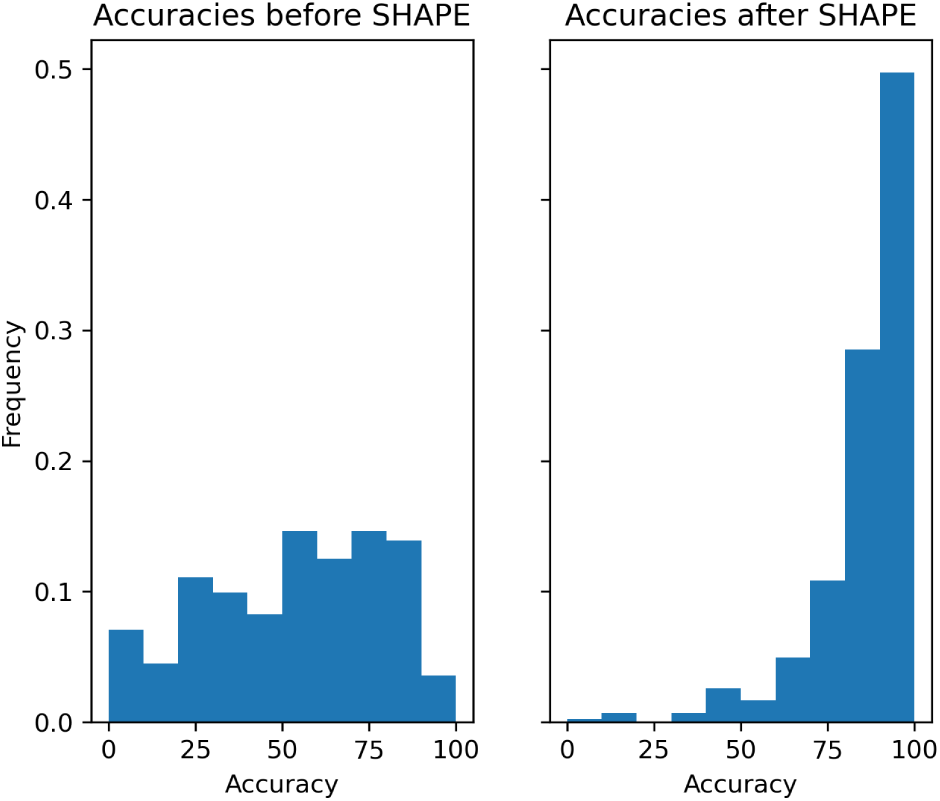
Accuracies before and after synthetic SHAPE for TestSetB. as an appropriate tool for this kind of task.

## 4 Results

In this section, we report the accuracy and feature importance of the different classifiers for well-prediction and directability. Our classifiers are based on the Random Forest Classification algorithm, which we selected

### 4.1 Classifying Well-Prediction

We built classifiers that take an RNA sequence as an input and then predict whether the secondary structure of the sequence is well-predicted (WP) or not (NWP) by the NNTM model, see Figures 8-9. We considered an RNA sequence as being WP if its NNTM accuracy (i.e., the MFE accuracy) is higher than an accuracy threshold. Otherwise, the sequence is classified as NWP. We varied the accuracy thresholds from 50% to 60%, these are given in the *y*-axis of Figures 8-9. At these thresholds, the original (unbalanced) Training and Test sets contain an approximately equal number of WP and NWP sequences. Thus, our balanced Training and Test sets contain nearly all of the sequences in the original (unbalanced) sets. These thresholds maximize the number of sequences available for training while ensuring the training set is balanced.

**Figure 8:**
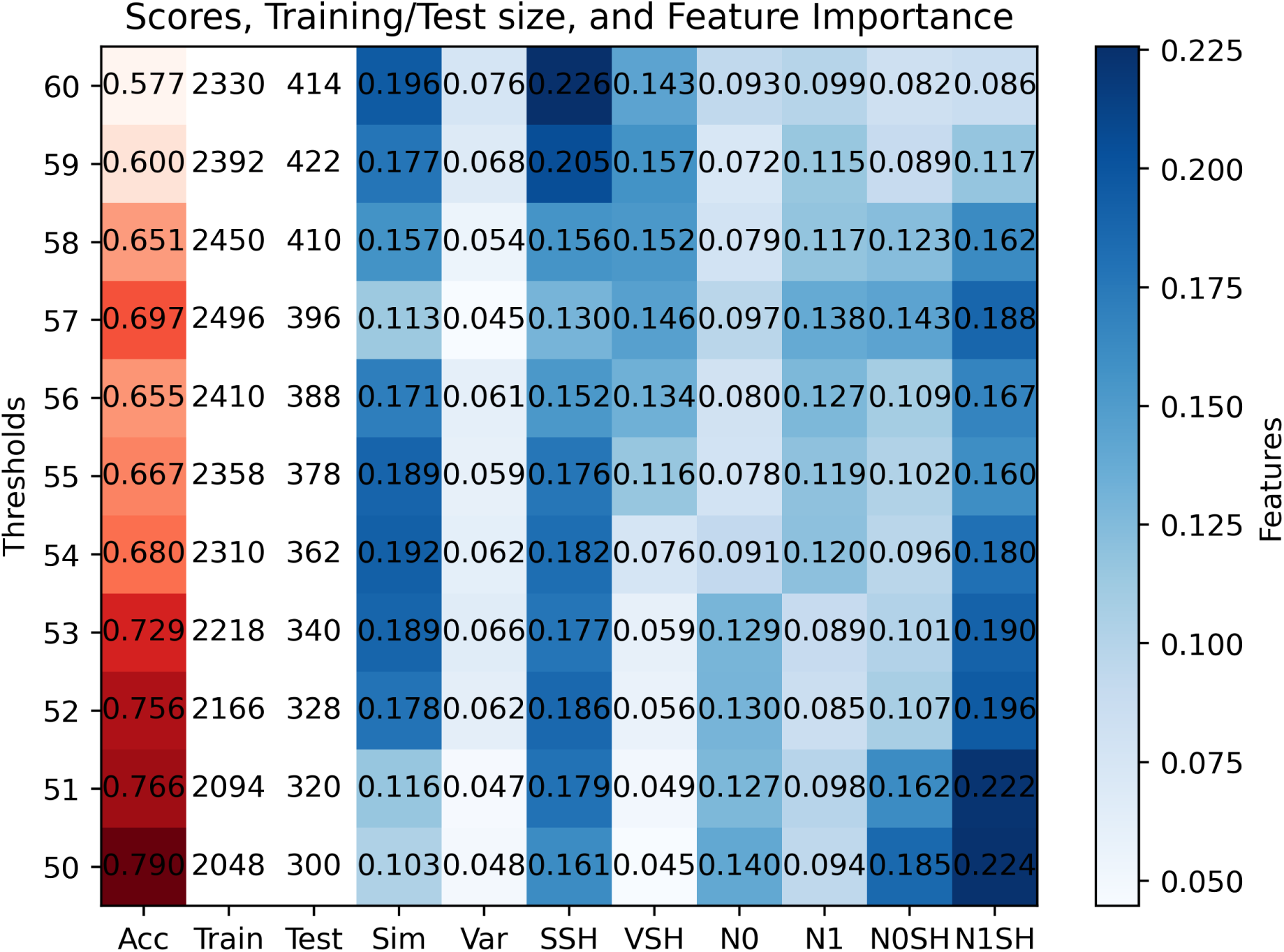
Classifying well-prediction on TestSetA. The classifiers were trained using sequences from Train-SetA and then tested using sequences from TestSetA. The first column (Acc) gives the accuracy of the classifier. The red shades correspond to the range of accuracies in the first column while the blue shades correspond to feature importance contributions in columns 4-12. The second (Train) and third (Test) columns give the number of RNA sequences in the Training/Test sets. Both Training and Test sets were balanced, that is, they contain equal number of sequences corresponding to each classification label.

**Figure 9:**
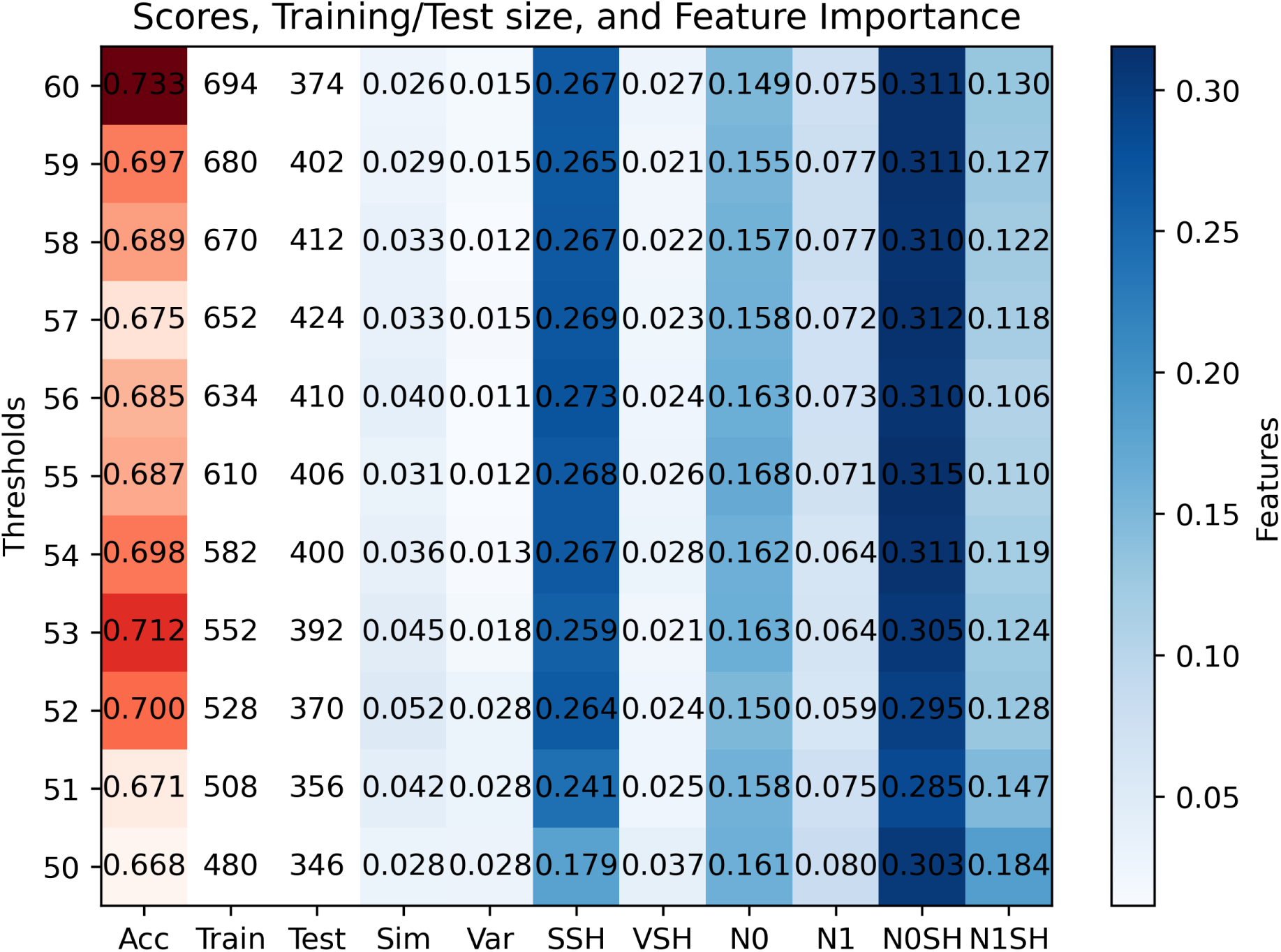
Classifying well-prediction on TestSetB. The classifiers were trained using sequences from TrainSetB and then tested using sequences from TestSetB. The first column (Acc) gives the accuracy of the classifier. The red shades correspond to the range of accuracies in the first column while the blue shades correspond to feature importance contributions in columns 4-12. The second (Train) and third (Test) columns give the number of RNA sequences in the Training/Test sets. Both Training and Test sets were balanced, that is, they contain equal number of sequences corresponding to each classification label.

Our classifiers were created in Python using the RandomForestClassifier implementation from SciKitLearn. Each RandomForestClassifier object contains the following hyperparameters: **n estimators** (the number of trees in the random forest), **max depth** (the maximum depth of a tree), and **random state** (determines the random number generator used). Unless stated otherwise, we set *n estimators*=1000, max depth=2, and random st=0. Classifiers were constructed using one of two training sets (TrainSetA and TrainSetB). Training sets were balanced to contain an equal number of WP and NWP sequences. We tested classifiers using data from one of two test sets (TestSetA and TestSetB) and obtained a **score** from Python, which is the percentage of sequences in the test set that were correctly identified as WP or NWP.

From Figure 8, which corresponds to classifiers trained in sequences in TrainSetA and tested in sequences in TestSetA, we see that for the task of well-prediction, the most important features are the similarities before SHAPE (Sim), the similarities after SHAPE (SSH), and the 1-nom after SHAPE (N1SH). And, in Figure 9, which corresponds to classifiers trained in sequences in TrainSetB and tested in sequences in TestSetB, we see that for the task of well-prediction, the most important features are the similarities after SHAPE (SSH) and the 0-norm after SHAPE (N0SH). The reason for the difference in feature importance between the classifiers trained in TrainSetA and the classifiers trained in TrainSetB is likely due to the composition of these two training sets of sequences. As noted in [3], TrainSetB and TestSetB contain dissimilar structures, while TrainSetA and TestSetA have similar structures. Thus, we expect the similarity feature (Sim) to be more important for classifiers trained on TrainSetA than for classifiers trained on TrainSetB. We also note that the 0-norm after SHAPE feature (N0SH) is more important for classifiers trained in TrainSetB than for classifiers trained in TrainSetA, indicating that the mechanisms for improving accuracy through the addition of auxiliary data have different effects on sequences of TrainSetA and TrainSetB. The effect of adding auxiliary information on the profile graphs of TrainSetB seems to be on the number of vertices (most likely reducing the number of competing alternates) while for the profile graphs of TrainSetA seems to be on the 1-dimensional holes (likely resolving conflicts). Finally, we note that the similarity after SHAPE (SSH) feature was important for both types of classifiers. This suggests that this feature captures the effect of the auxiliary data on the competing substructures, making it useful for this type of classification task.

### 4.2 Classifying directability

We built classifiers that take an RNA sequence as input and predict whether the sequence is directable (DIR) or not directable (ND). For these classifiers, we considered an RNA sequence as directable if the improvement in accuracy after adding auxiliary information is higher than a directability threshold. Otherwise, the sequence is classified as not directable. We varied the directability thresholds from 25% to 50%, these are given in the *y*-axis of Figures 10 and 11. At threshold=35, our original (2503 sequences) TrainSetA has an approximately equal number of DIR and ND sequences, such that our balanced TrainSetA (2476) contains nearly all of the original set. Thus, we selected directability thresholds around 35 to maximize the size of training. Additionally, we observed a notable increase in accuracy for TrainSetA as the directability threshold increase, thus, we varied the directablity threshold up to threshold=50 to capture this trend.

**Figure 10:**
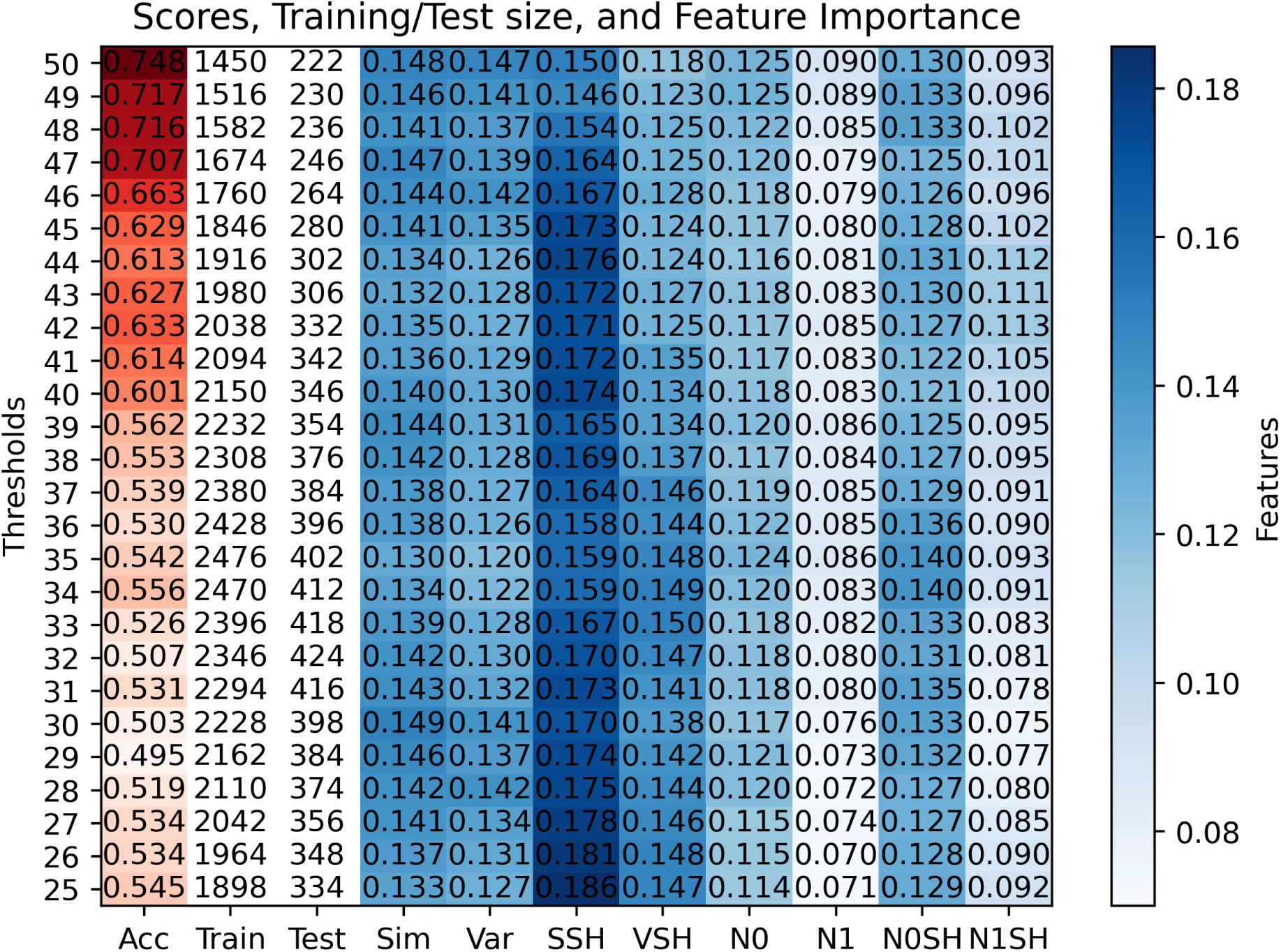
Classifying directability on TestSetA. The classifiers were trained using sequences from TrainSetA and then tested using sequences from TestSetA. The first column (Acc) gives the accuracy of the classifier. The red shades correspond to the range of accuracies in the first column while the blue shades correspond to feature importance contributions in columns 4-12. The second (Train) and third (Test) columns give the number of RNA sequences in the Training/Test sets. Both Training and Test sets were balanced, that is, they contain equal number of sequences corresponding to each classification label.

**Figure 11:**
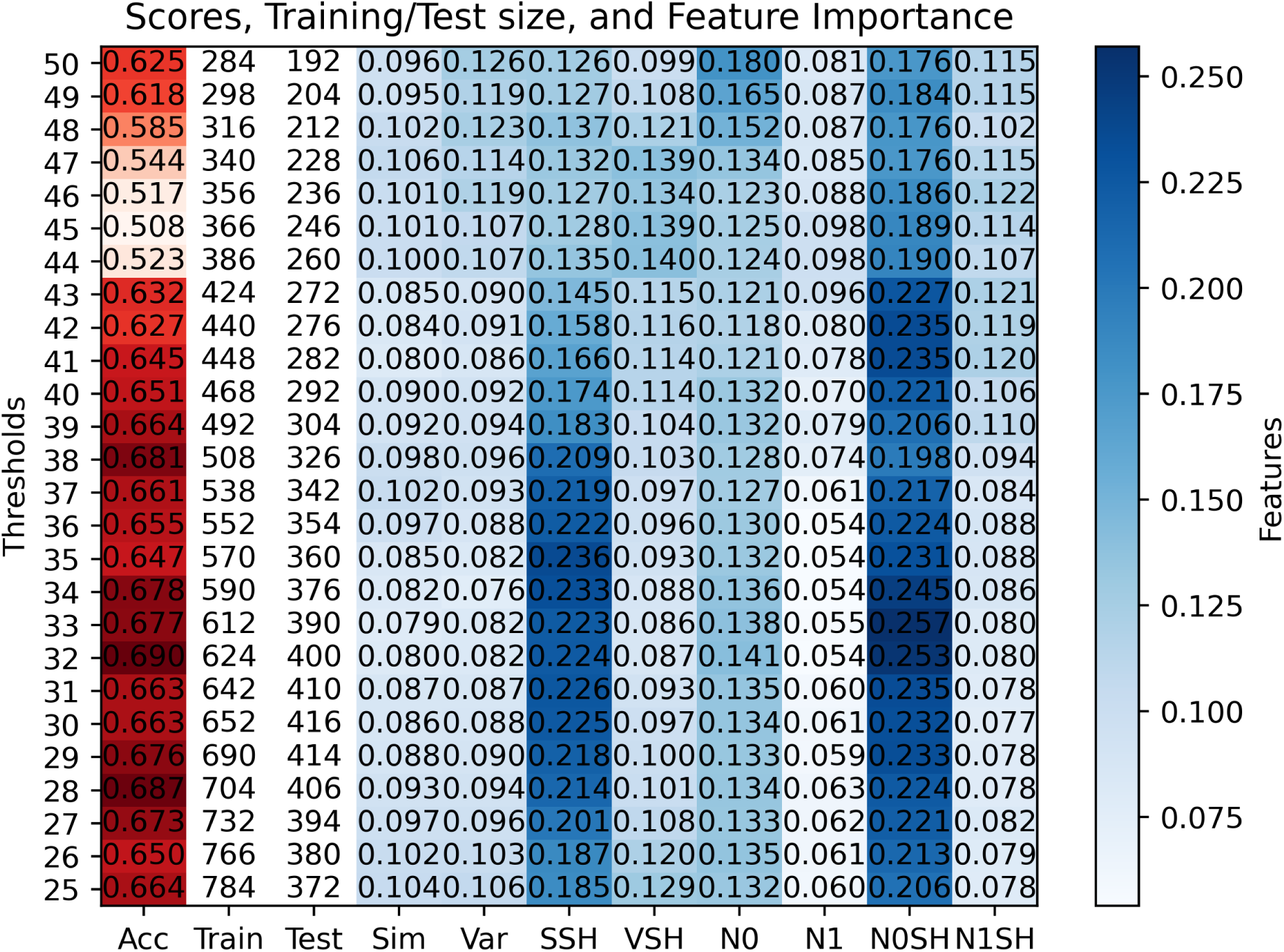
Classifying directability on TestSetB. The classifiers were trained using sequences from TrainSetB and then tested using sequences from TestSetB. The first column (Acc) gives the accuracy of the classifier. The red shades correspond to the range of accuracies in the first column while the blue shades correspond to feature importance contributions in columns 4-12. The second (Train) and third (Test) columns give the number of RNA sequences in the Training/Test sets. Both Training and Test sets were balanced, that is, they contain equal number of sequences corresponding to each classification label.

From Figure 10, which corresponds to classifiers trained in sequences in TrainSetA and tested in sequences in TestSetA, we see that for the task of directability, the most important feature is the similarity after SHAPE (SSH). And, from Figure 11, which corresponds to classifiers trained in sequences in TrainSetB and tested in sequence in TestSetB, we see that for directability, the most important features are the similarity after SHAPE (SSH) and the 0-norm after SHAPE (N0SH). Similar to the case with the well-prediction classifiers, the difference in feature importance between classifiers trained in TrainSetA and classifiers trained in TrainSetB is likely due to differences in the composition of these two training sets of sequences. We note that the sequences in TrainSetB seem harder to direct (compared to the sequences in TrainSetA) as shown in Figures 4-5. Additionally, we observe that the N0SH feature captures the changes in size of the profile graphs, therefore it correlates with the number of competing profiles. Therefore, the fact that the N0SH feature is more important for classifiers trained in TrainSetB than for classifiers trained in TrainSetA (for both well-prediction and directability) highlights the differences in the effects of auxiliary information for directability when considering training sets with dissimilar sequences. Finally, as was the case for well-prediction, the similarity after SHAPE feature (SSH) was important for directability classifiers, indicating that this feature captures the effect of the auxiliary data on the competing substructures, making it useful for this type of classification task.

### 4.3 Predicting both accuracy and directability

Here we consider a case where we first predict well-prediction and then the directability of a sequence. Building this pipeline of classifiers was motivated by the idea that training on subclasses of sequences with similar properties (e.g., being already well-predicted or directable) improved the accuracy of the classifiers. Thus, we trained classifiers on sequences that are already well-predicted and on sequences that are not well-predicted. The pipelines in Figures 12-13 consider three classifiers. The first classifier is for well-prediction. This is similar to the classifier that we considered in Section 4.1. The other two classifiers (classifiers 2 and 3) are similar to those considered in Section 4.2, but these classifiers were trained only on sequences that are already well-predicted (classifier 2) and on sequences that are not well-predicted (classifier 3).

**Figure 12:**
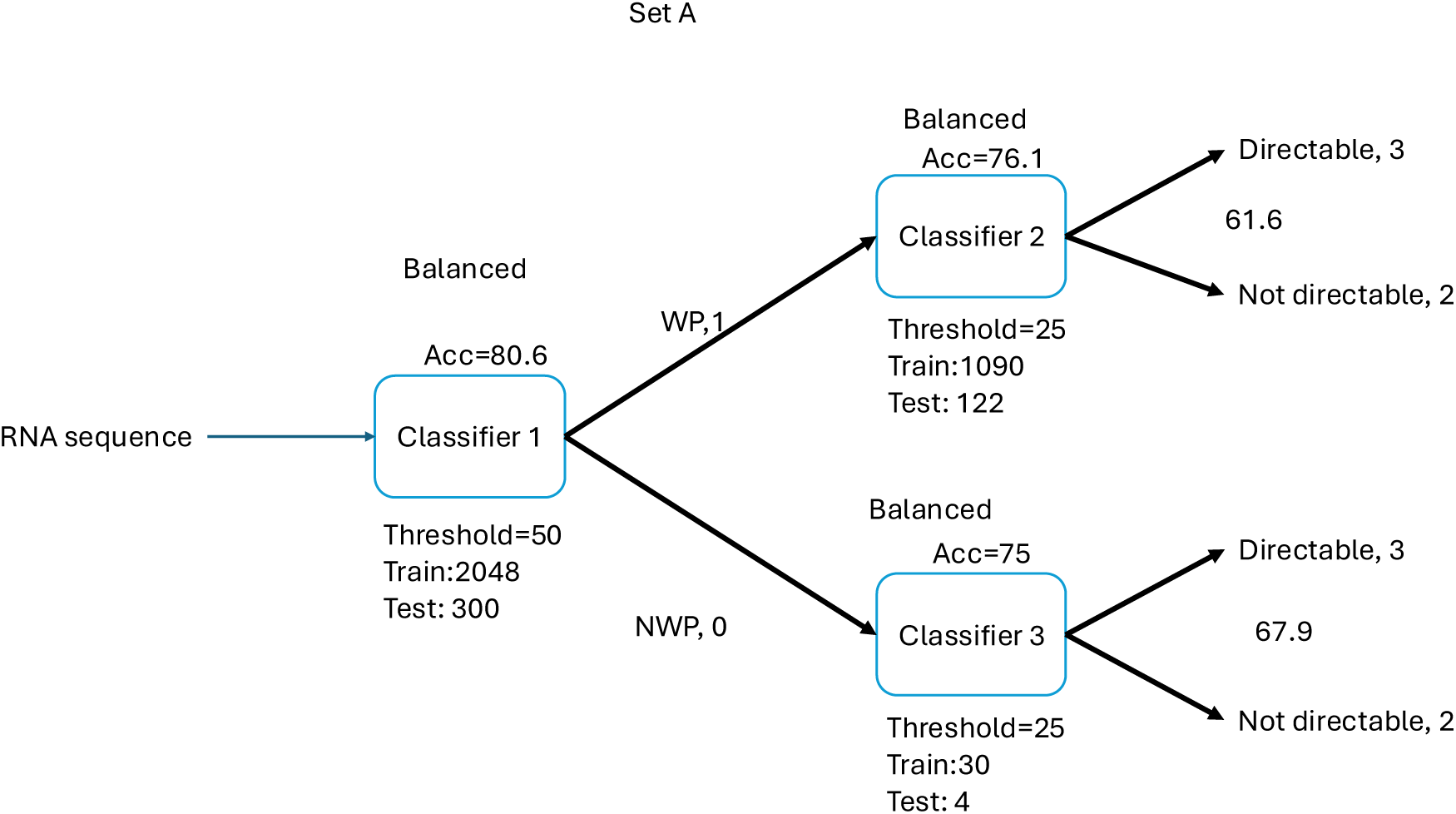
Classifying well-prediction and then directability on TestSetA. The classifiers were trained using sequences from TrainSetA and then tested using sequences from TestSetA. The labels for Classifier 1 are well-predicted (WP, 1) and not-well-predicted (NWP, 0) while the labels for Classifiers 2 and 3 are directable (DIR, 3) and not-directable (ND, 2). The training sets for each classifier have been balanced with equal number of sequences for each label.

**Figure 13:**
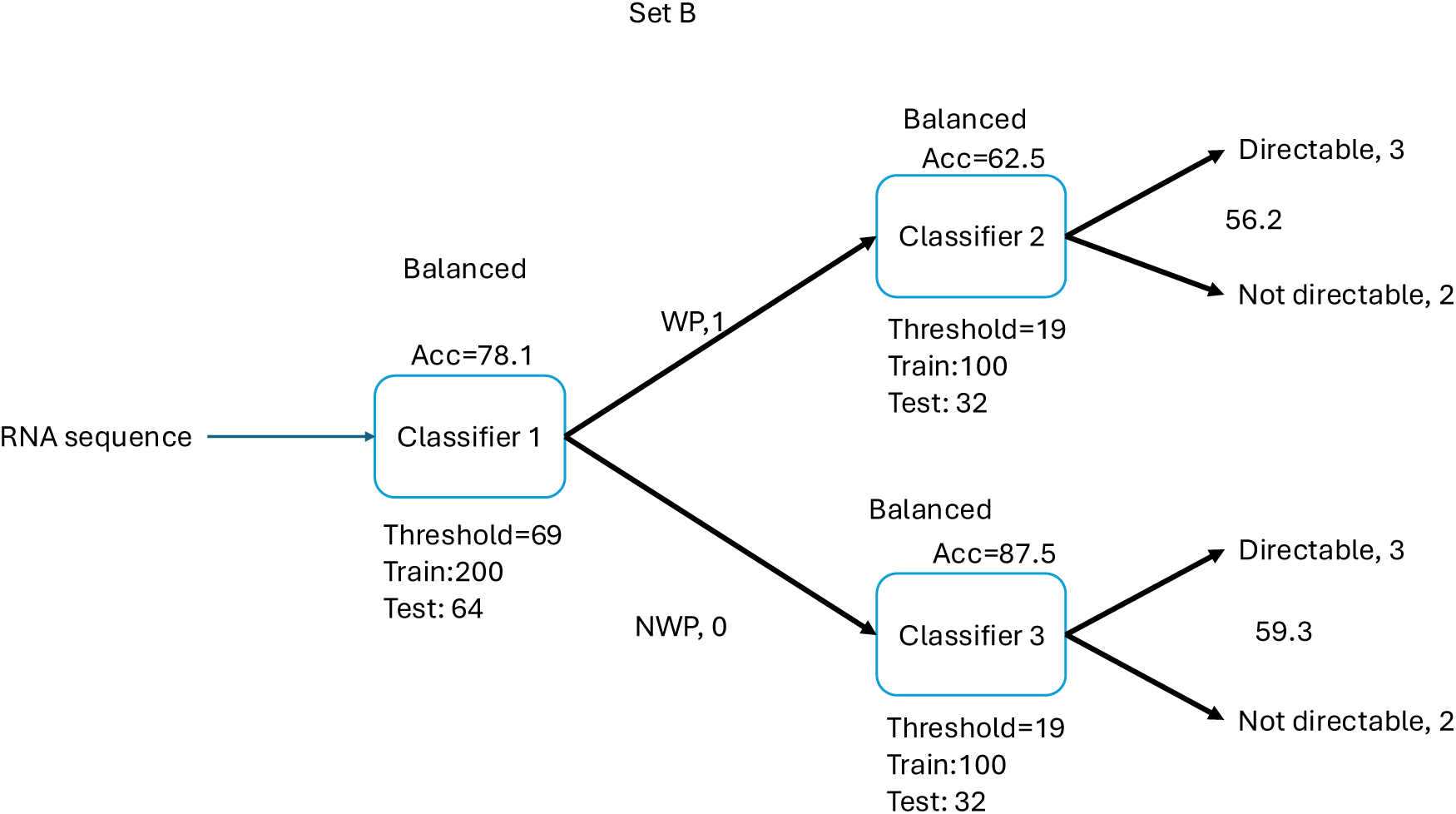
Classifying well-prediction and then directability on TestSetB. The classifiers were trained using sequences from TrainSetB and then tested using sequences from TestSetB. The labels for Classifier 1 are well-predicted (WP, 1) and not-well-predicted (NWP, 0) while the labels for Classifiers 2 and 3 are directable (DIR, 3) and not-directable (ND, 2). The training sets for each classifier have been balanced with equal number of sequences for each label.

For example, in Figure 12, we first split the training set into well-predicted (WP=1) and not-well-predicted (NWP=0) sequences. We then classify the well-predicted sequences according to directability using Classifier 2 into directable (DIR=3) and not-directable (ND=2). Similarly, we classify the not-well-predicted sequences according to directability using Classifier 3 into directable (DIR=3) and not-directable (ND=2). This establishes a two-step pipeline to classify sequences first according to well-prediction and then directability. To test this pipeline, we considered a balanced test set of n=300 sequences (150 WP, 150 NWP). Note that this is the test set in which Classifier 1 yields the greatest accuracy. First, we use Classifier 1 to classify each sequence in the test set as well-predicted (WP=1) or not-well-predicted (NWP=0). Next, we only consider those sequences which were predicted as WP by Classifier 1 and classify these according to directability with Classifier 2. This yields an accuracy of 61.6 percent. Similarly, we consider those sequences which were predicted as NWP by Classifier 1 and classify these according to directability using Classifier 3. This gives us an accuracy of 67.9 percent.

The classifiers in Figure 12 were trained in TrainSetA and tested on sequences in TestSetA. Similarly, the classifiers in Figure 13 were trained in TrainSetB and tested on sequences in TestSetB where we balanced the training and test sets to have an equal number of sequences for each label. In both cases, we balanced the training and test sets so that there is an equal number of sequences for each classification label.

Here we note that restricting the training sets to sequences with similar properties (e.g., only well-predicted sequences) allows us to achieve higher accuracies as shown in the accuracies of the middle subclassifiers in Figures 12-13. Although, we achieved higher accuracies on the subclassifiers of the pipelines of Figures 12-13, the overall accuracy of each pipeline was lower than the direct classifier for directability.

## 5 Discussions

In this work, we have used synthetically generated auxiliary data to study the role of structural features in determining accuracy improvements via auxiliary information. Although experimentally generated auxiliary data (i.e., SHAPE data [12]) are available for different sets of RNA sequences, we are not aware of experimental SHAPE data for the set of sequences that we have used in this study. Therefore, we acknowledge that the validation of our results using experimentally generated SHAPE data might be a limitation of this investigation. However, synthetically generated auxiliary information have been shown to achieve similar accuracy improvements to the experimental data [14] and it has been used to evaluate the extend of accuracy improvements in RNA secondary structure prediction the past [14, 15, 16].

For our random forest classifiers, to avoid bias in our results we have used balanced training/test sets. That is, the training/test sets were balanced by containing an equal number of sequences corresponding to each classification label. We acknowledge that such balance may not occur in nature and that might impact the generalizability of our results. However, if working with different RNA datasets, we still recommend working with balanced training sets as a good practice to prevent bias in subsequent results.

We have used a range thresholds between 50% to 60% for the well-prediction task to allow enough room for directability. Likewise, we considered a range of threshold between 25% to 50% for the directability classification as these amounts of increase in accuracy would allow to the correct prediction of a relevant region and/or motif in the secondary structure of an RNA sequence. We also note that at these thresholds, the original (unbalanced) Training and Test sets contain an approximately equal number of WP and NWP sequences. Thus, our balanced Training and Test sets contain nearly all of the sequences in the original (unbalanced) sets. These thresholds maximize the number of sequences available for training while ensuring the training set is balanced.

## 6 Conclusions

In this paper, we considered features based on suboptimal formations competing with the minimum-free energy formation and investigated their role in determining well-prediction and directability. First, we introduced a similarity measure among competing substructures called profiles which was used to generate features for Random forest classifiers. Second, we presented an *n*-dimensional representation of the profiles which allows the use of topological data analysis (specifically, persistence landscapes) to obtain different metrics that represent topological features. Third, we built random forest classifiers using these similarities and topological features. Fourth, we show that the similarity features are more important for classifiers trained on sequences with similar structures, while the topological features are more important for classifiers trained on sequences with dissimilar structures. Finally, we performed extensive testing on two sets of RNA sequences where we studied the sensitivity of the classification accuracy and their feature importance. Overall, we identified the similarity after SHAPE (SSH) and the 0-norm after SHAPE (N0SH) are the most important features for directability and well-prediction. We also showed that one can achieve higher accuracies by restricting the training sets to sequences with similar properties such as training only on sequences that are well-predicted.

## 7 Acknowledgments

Part of this work was supported by a pilot grant (to D.M., L.S.G, and L.R.) from the University of Kentucky AI/ML Hub initiative. D.M. was partially supported by a Collaboration grant (# 850896) from the Simons Foundation and a grant (NSF: # 2424633) from the National Science Foundation. D.N. was partially supported by the National Science Foundation (NSF: # 2516126, # 2151802, and # 2534947). M.W. was partially supported by the National Science Foundation (NSF:#2424633) and the National Institute of Health (NIH: R01-AI135128).

## 8 Code and Data Availability

Python code in Jupyter notebooks that we used for training Random Forest classifiers are available on a GitHub repository at https://github.com/LoganRose25/RNA-project. We also provide the required data for training and testing these classifiers in csv files in the same repository. For persistence landscape computations done in R, the TDA-tools package is required. This R package and installation instructions can be found here: https://github.com/jjbouza/tda-tools. All other data supporting the findings of this study are available within the cited papers and their supplementary information.

## 9 Author Contributions

Conceptualization: D.M., D.N., and L.S.G.; Methodology: L.R., M.W., and D.M., Software: L.R. and M.W.; Formal analysis: L.R., M.W., and D.M.; Supervision: D.M.; Writing—original draft: D.M.; Writing—review and editing: L.R., L.S.G., D.N., M.W., and D.M.

